# Natural image reconstruction from brain waves: a novel visual BCI system with native feedback

**DOI:** 10.1101/787101

**Authors:** Grigory Rashkov, Anatoly Bobe, Dmitry Fastovets, Maria Komarova

**Affiliations:** Neurobotics LLC, Moscow, Russian Federation; Neuroassistive Technologies LLC, Moscow, Russian Federation; Moscow Institute of Physics and Technology, Dolgoprudny, Moscow Region, Russian Federation

## Abstract

Here we hypothesize that observing the visual stimuli of different categories trigger distinct brain states that can be decoded from noninvasive EEG recordings. We introduce an effective closed-loop BCI system that reconstructs the observed or imagined stimuli images from the co-occurring brain wave parameters. The reconstructed images are presented to the subject as a visual feedback. The developed system is applicable to training BCI-naïve subjects because of the user-friendly and intuitive way the visual patterns are employed to modify the brain states.

## 1. Introduction and related work

Currently, the usage of EEG-based BCIs in assistive and rehabilitation devices mostly comes down to the following scenarios:

1) using synchronous BCIs (based on event-related potential registration, e.g. P300) for making discrete selections;
2) using asynchronous BCIs based on motor imagery potential or concentration/decocentration-driven mental states for issuing voluntary controlling commands.

Both scenarios have some advantages which are, unfortunately, overweighed with severe limitations that hinder implementations of BCI technology in real-world tasks. Thus, in synchronous BCI paradigms, a wide variety of stimuli, including visual categories, can be utilized to explore and measure the evoked responses of a particular subject [1]. However, the whole set of stimuli has to be successively presented to the subject each time to determine his intention, which makes such approach inconvenient for the applications requiring fast, real-time control of an external device. Motor-imagery or other asynchronous BCIs do not require any external stimuli presentation, which allows a subject to produce voluntary mental commands at his own wish. At the same time, the ability of different subjects to perform various mental tasks is variable and depends on their personal physiological parameters and experience [2]. A typical asynchronous BCI scenario requires an unexperienced subject to undergo a long-time training routine to master the control of at least 2 or 3 mental states. The practical instructions on how to perform abstract mental tasks are often unclear to novice BCI operators, which adds to overall complexity of the procedure. As a consequence, different studies show that classification rates are highly inconsistent even for similar paradigms [3–5].

A solution could be to join the advantages of the two scenarios by exploring the effect of continuous stimuli presentation on brain wave patterns. The decoded long-term evoked responses, if any, could then be treated as subject-specific mental states which could potentially be triggered by a particular stimuli imagination. Bobrov et.al. showed that perception or imagination of static images of different categories provide distinct EEG patterns which can be used for BCI [6]. Another suggestion is to use natural movies of different objects as attention-capturing continuous stimuli. This kind of stimulation has been already reported for fMRI research [7]. Grigoryan et.al. have recently showed the positive effect of mobile visual stimuli in EEG-based BCIs as well [8].

Another essential part of most BCI protocols is maintaining a neurofeedback loop for the subject. Many articles have shown that the self-regulation effects achieved through this approach could facilitate learning of mental states [9–13]. Ivanitsky et.al. figured out an important role of BCI feedback even for mastering complex cognitive tasks [14]. Ideally, a feedback should give a subject sufficient information about his progress, but not distract him from performing the mental task itself, i.e. being native in terms of the BCI paradigm. In recent works Horikawa et.al. [15, 16] propose a model for natural image reconstruction from fMRI brain signal data recorded from a subject while he observes the original images. There have been similar reports on similar EEG-based reconstructions [17], but the reliability of the studies has met serious controversy [18].

Considering the conditions outlined above, one can reasonably suggest that closed-loop asynchronous BCIs with adaptively modifiable set of mental states and a native type of feedback could outperform the other BCI approaches. In this article, we introduce a novel BCI paradigm that meets these requirements. Our protocol features the visual-based cognitive test for individual stimuli set selection as well as state-of-art deep learning based image reconstruction model for native feedback presentation.

## 2. Methods

For our research we set two major objectives:

- Exploring the continuous effect of visual stimuli content on rhythmical structure of subject’s brain activity.
- Developing a model for mapping the EEG features extracted for a given category of observed stimuli back into the natural image space of the same category.

### 2.1. Subjects

The human protocol was approved by the local Institutional Ethical Committee (#5 of 18.05.2018). We recruited 17 healthy subjects with no history of neurology diseases, 11 males, 6 females, all right-handed. The subjects’ age ranged from 18 to 33 years (mean age: 22). The subjects were informed about the experimental protocol and signed the informed consent form.

### 2.2. EEG recordings

The EEG recording equipment included a 128-channel EEG cap and NVX-136 amplifier developed by Medical Computer Systems Ltd. (Moscow, Zelenograd). The conductive gel (Unimax gel) was injected into each electrode. Each subject was seated in a comfortable position at a distance approximately 0.6m from an LCD computer monitor. The EEG signals were recorded at a sampling rate of 500 Hz with NeoRec software (developed by Medical Computer Systems Ltd.). The recorded signals were filtered using a band pass filter with a bandwidth of 1-35Hz.

### 2.3. Visual stimuli

In this study, we used video clips of different objects as stimuli. We assumed that observing the videos rather than the static images would keep the subjects motivated during the experiment, making an extra mental task unnecessary. We selected following stimuli object categories, which could potentially affect the subjects’ mental state by inducing relaxation, concentration on particular items, or imposing stress (the examples are shown on Figure 1):

- A - abstract geometric shapes or fractals (visual illusions);
- W - natural waterfalls;
- HF - human faces with different emotions;
- GM - Goldberg mechanisms (mechanical devices with a large number of elements triggering each other);
- E - extreme sports (first-person videos of high speed motion activities, some ending with accidents).

**Figure 1.**
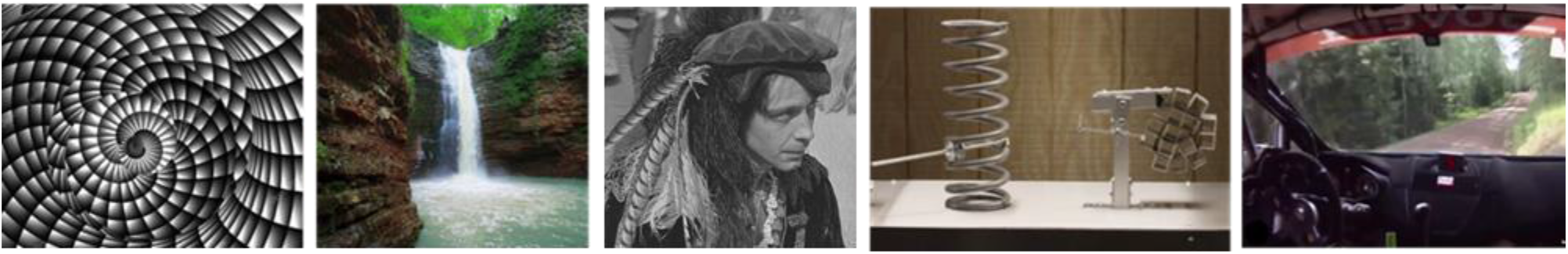
Examples of video frames from each of the stimuli categories

The experiment consisted of two sessions with a short break of 5-10 minutes between them. During each session, a subject was asked to watch a 21-minute video sequence comprised of 117 randomly mixed video clips. The duration of a single clip was between 6 and 10 seconds, and the “black screen” transitions of 1-3 seconds were inserted between the consecutive clips.

The video sequence contained 25 clips of each category except A, where only 17 fragments were present due to low variability of the category and the fatigue effect caused by such videos. There were no identical clips within a category, as well as no montage transitions within a single clip to avoid the occurrence of parasite ERPs. The onsets of the video clips were precisely synchronized with EEG signal using MCS VGASens photo sensor.

### 2.4. Feature extraction and classification

The EEG data was epoched into time segments corresponding to each single clip observation, and each time segment was split into 3-second time windows with 2/3 overlap.

Independent component analysis (ICA) matrix was calculated using the training data only. The muscle and ocular artifact rejection procedure included manual selection and removing of the independent components which contained specific patterns from blinks, eye saccades, motions and pulse. All of the following signal processing stages were performed within ICA space as well. We used fast Fourier transform to extract the spectral features for each component remaining after artifact removal. As dense electrode recordings produce a way excessive feature dimensionality, we considered a scoring procedure for feature space compression. The average power spectrum values were obtained for each *k*-th data sample of each *n*-th component using a sliding frequency window *w* of 3 Hz:

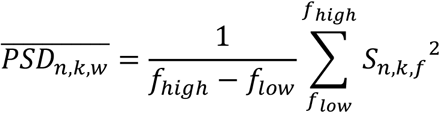

A set of simple threshold classifiers was created separately for each of the PSD values. The features were scored according to these classifiers performance on the training set. As each classifier gave an estimation of predictive power related to a particular frequency band of a particular ICA component, the components with the best overall feature scores were selected and the most informative feature bands were specified for each of the selected components.

The dimensionality of the feature vectors was then further reduced to a fixed value of 20 using principal component analysis (PCA) transform. The feature classification was performed using linear discriminant analysis (LDA).

## 3. Model evaluation

In order to simulate the real-world procedure of subject training, we used one session for training the model and another one for performance validation. We also switched the training and validation sessions and averaged the results of this cross-validation for final estimation.

Table 2 presents the rates for pairwise category classification task. For each pair of classes, we balanced the number of features by randomly dropping some samples from the class that contained more data. The significance level for each binary classifier was estimated with a one-tailed binomial test. As we used overlapping windows for evaluation, we used the corresponding number of actually independent trials for setting the unbiased classifier confidence rate. The actual and recalculated number of samples and the corresponding confidence rates for non-random hypothesis acceptance are shown in Table 1.

**Table 1.** Confidence rates for different binary classifiers

| | Number of<br>oversampled features<br>(for classifier) | Number of independent trials<br>(for binomial test) | Non-random<br>classification rate<br>($p=0.95$) |
| --- | --- | --- | --- |
| For “A-W”, “A-HF”,<br>“A-GM”, A-E” | 320 | 68 | 0.61 |
| For Others | 614 | 130 | 0.57 |

**Table 2.** Classification results: averaged accuracy values.

| Subj | A-W | A-HF | A-GM | A-E | W-HF | W-GM | W-E | HF-GM | HF-E | GM-E |
| --- | --- | --- | --- | --- | --- | --- | --- | --- | --- | --- |
| 1 |  |  |  |  | 0.73 | 0.83 | 0.64 | 0.85 | 0.79 | 0.74 |
| 2 | 0.72 |  | 0.83 |  |  | 0.66 |  | 0.82 | 0.57 | 0.74 |
| 3 |  |  | 0.76 | 61 | 0.68 | 0.77 | 0.68 | 0.83 | 0.72 | 0.82 |
| 4 |  | 0.73 | 0.73 |  | 0.59 | 0.78 | 0.69 | 0.85 | 0.85 | 0.67 |
| 5 |  |  | 0.85 |  | 0.64 | 0.77 | 0.62 | 0.89 | 0.86 | 0.77 |
| 6 |  |  | 0.62 |  |  |  | 0.72 | 0.77 | 0.73 | 0.61 |
| 7 |  |  |  |  |  | 0.69 | 0.64 | 0.78 | 0.65 | 0.70 |
| 8 | 0.73 |  | 0.83 |  | 0.75 | 0.78 | 0.69 | 0.83 | 0.72 | 0.78 |
| 9 | 0.75 |  | 0.67 | 61 | 0.60 | 0.57 | 0.68 | 0.59 | 0.64 | 0.73 |
| 10 |  | 0.77 |  |  | 0.71 | 0.63 | 0.63 | 0.85 | 0.73 | 0.65 |
| 11 |  | 0.75 | 0.81 |  |  | 0.82 | 81 | 0.90 | 0.85 | 0.76 |
| 12 |  |  |  |  | 0.58 | 0.60 |  |  | 0.62 |  |
| 13 | 0.76 | 0.71 | 0.64 |  |  | 0.81 | 0.75 | 0.85 | 0.79 | 0.75 |
| 14 | 0.82 | 0.75 | 0.89 | 80 | 0.59 | 0.77 | 0.83 | 0.87 | 0.87 | 0.81 |
| 15 |  | 0.63 | 0.78 | 61 | 0.66 | 0.84 | 0.85 | 0.89 | 0.87 | 0.76 |
| 16 |  |  | 0.78 | 61 |  | 0.86 | 0.83 | 0.92 | 0.88 | 0.73 |
| 17 | 0.85 | 0.62 | 0.83 | 60 | 0.73 | 0.63 | 0.60 | 0.80 | 0.75 | 0.67 |
| $P_{nr}$ | 0.35 | 0.41 | 0.76 | 0.35 | 0.65 | 0.94 | 0.88 | 0.94 | 1.00 | 0.94 |
| $P_{rec}$ | 0.772 | 0.709 | 0.771 | 0.640 | 0.660 | 0.738 | 0.711 | 0.831 | 0.758 | 0.731 |

The classification accuracy scores below the confidence rates are marked as dark gray cells in Table 2. The overall results were evaluated using two metrics: the probability for the given binary classification to be non-random for a subject (*P*_*nr*_) and average classification accuracy among the subjects with non-random classification rate (*P*_*rec*_). 72.3% classifications proved to be at least non-random, with average score of 74.0% among them. We decided to withdraw the “abstract shape” category from further research due to generally mediocre results for our subjects, thus reaching 89.2% of non-random classifications with average score of 81.6% for the remaining 4 categories. We also trained a 4-class classifier for each subject, as well as for 5 top performing subjects (#1, 3, 11, 15, 16) and formed an averaged confusion matrix for the experiment (Table 3).

**Table 3.**
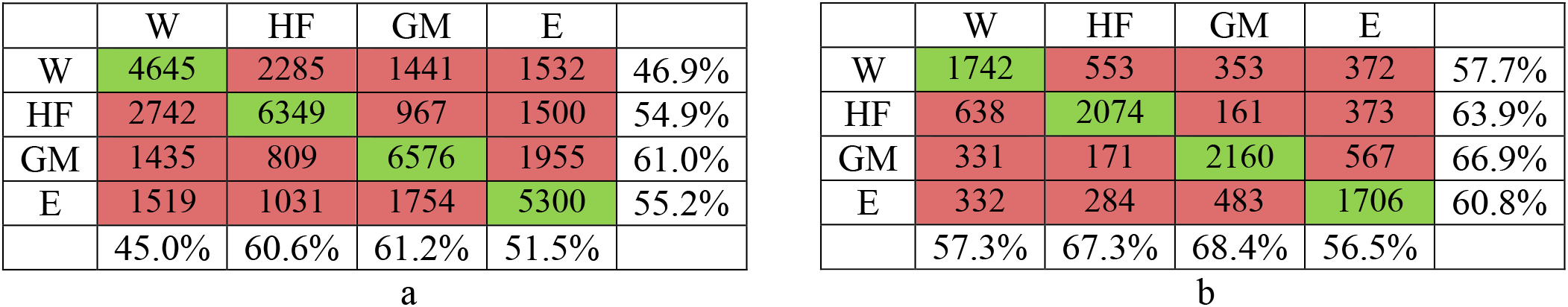
4-class confusion matrix for all subjects (a) and for top 5 subjects (b)

The examples of the of top-scored ICA components topographies (for 4-class classification task) are shown on Figure 2.

**Figure 2.**
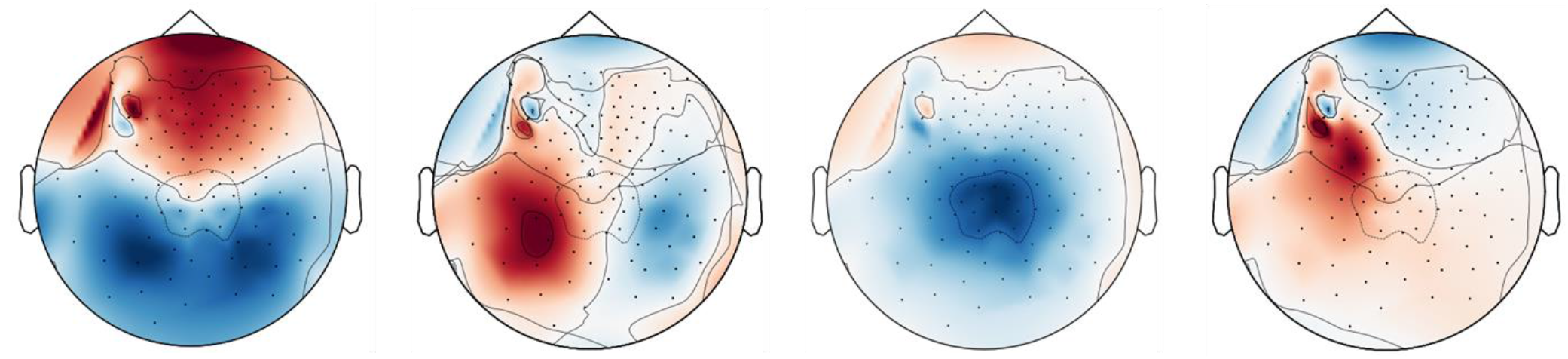
ICA topographies for top scoring ICA components (subject #15, 4-class model)

Overall, the model evaluation results show that some visual stimuli categories seem to be generally useful for BCI purposes among most of the subjects, and some other can be individually picked up or rejected depending on the results of the proposed cognitive test. Within each category, an exemplar-level analysis can be carried out to select the most effective stimuli for a particular subject.

## 4. Native neurofeedback model

The general idea of the proposed visual feedback is to present the BCI classifier predictions in form of natural images, which should be as close to the actually observed (or imagined) visual stimuli as possible. This would boost the subject’s imagination efforts by giving him an illusion of his thoughts being visualized and at the same time would not distract him with any side-triggers. A basic approach would be simply to show some image samples from the stimuli dataset according to the category predicted by the classifier. A strong drawback in this solution is that weak or uncertain predictions would be either strictly mapped to one of the categories or cause excessive image switching, both ways much confusing to the subject. We hypothesized that a proper visual feedback should satisfy the following criteria:

- dynamically represent the brain wave decoding process
- retain at least the general content of the reconstructed object so that a subject can always recognize the decoded category;
- avoid quick or sudden shifting between the category types even if the signal becomes noisy or inconsistent;
- represent classifier uncertainty in adequate form of “uncertain” image

In order to keep in terms with these requirements we developed a deep-learning based visualization model. The general scheme of the model is shown on Figure 3. The 20-dimensional EEG feature vector obtained after dimension reduction stage (see “*Feature extraction and classification*” section above) is mapped into the latent space of a pre-trained image autoencoder, which is capable of reconstructing natural images of several pre-learnt categories. An image decoding model is independent of any neurophysiological data and can be trained beforehand considering just a set of stimuli images. A feature mapping model is trained separately as it requires both EEG feature bank and a trained image decoder at use. In the following sections we explain the model training procedure in more detail.

**Figure 3.**
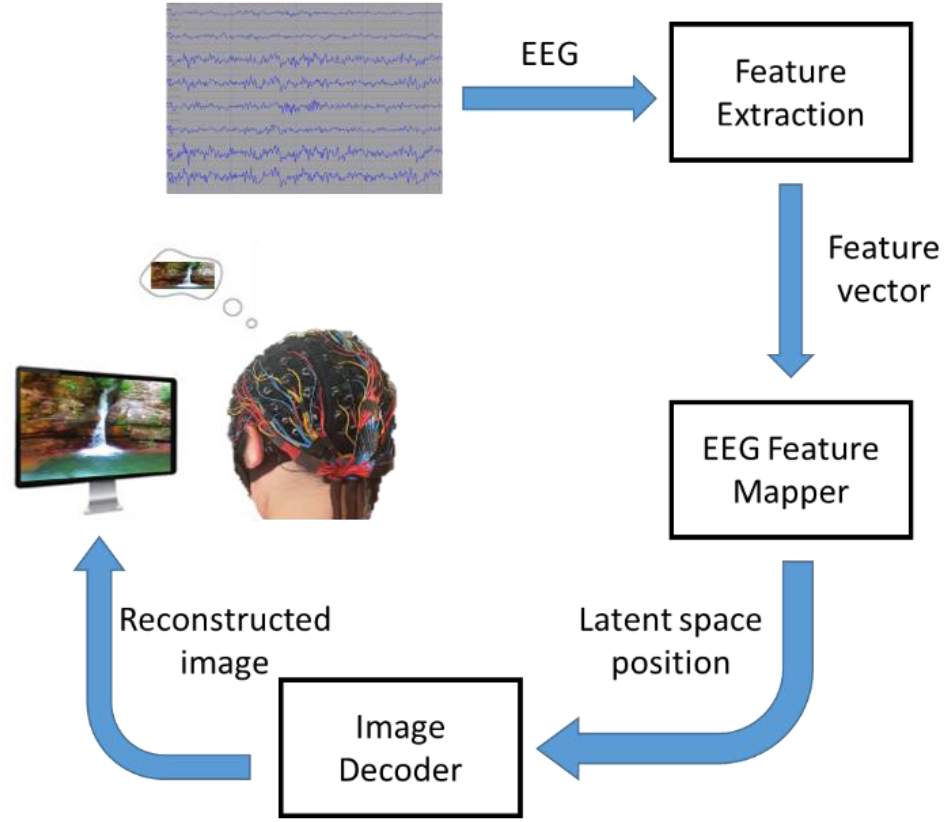
General scheme of a neurofeedback model

### 4.1. Image Decoder

An image decoder (ID) is a part of image-to-image convolutional autoencoder model. The encoder part is based on a pre-trained VGG-11 model [19]. The decoder part is composed of a fully-connected input layer for dimension enhancement, followed by 5 deconvolution blocks, each one containing a deconvolutional layer followed by rectifier linear unit (ReLU) activation. The final deconvolutional block contains hyperbolical tangent activation layer. A decoder produces color images of 192×192×3 dimensionality (see Figure 4a).

**Figure 4.**
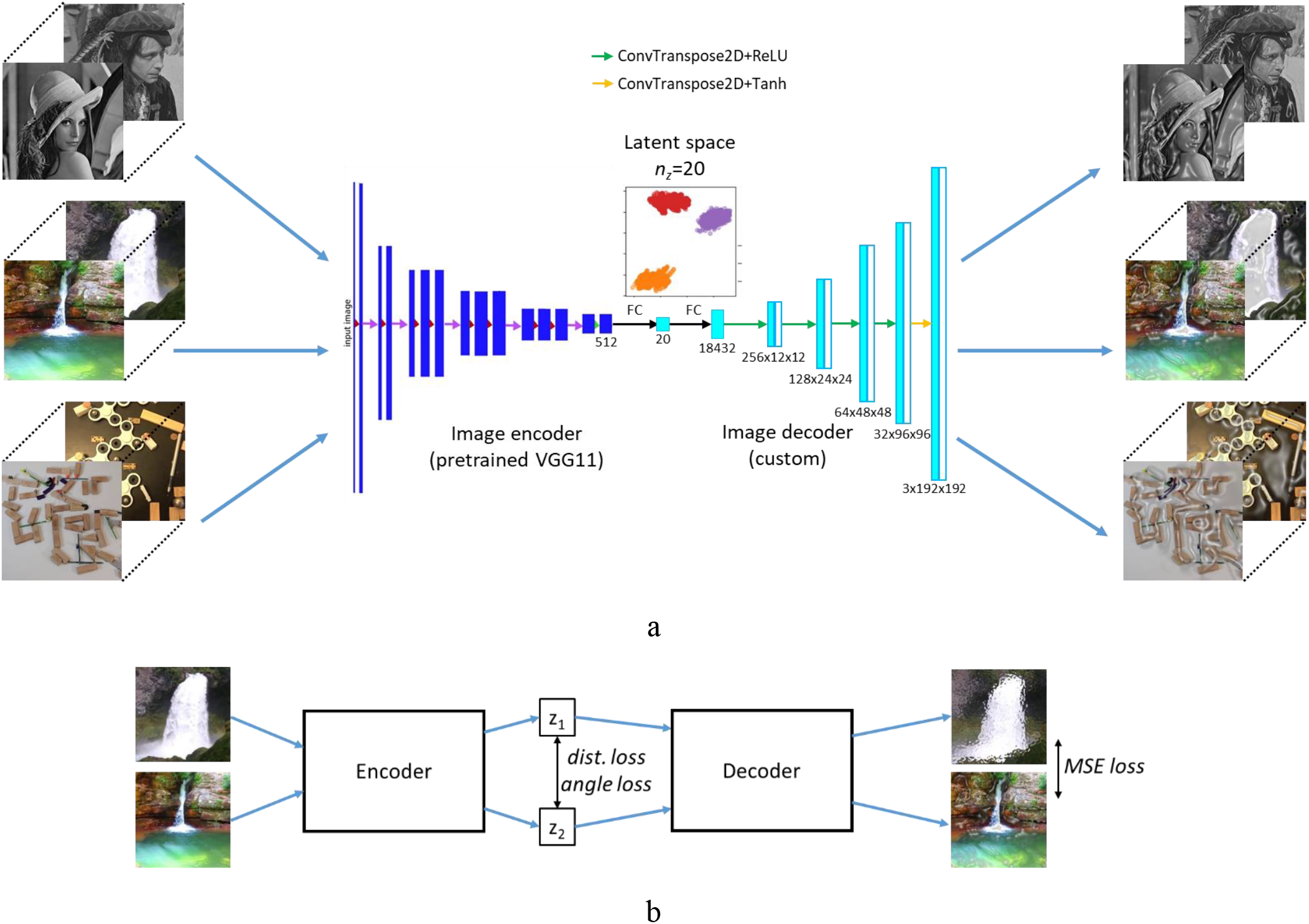
Image decoder. a) Model structure; b) Training routine

Apart from image reconstruction, we suggest our decoder model to have a specific distribution of its latent space. We handled this problem by introducing a training procedure shown on Figure 4b. The encoder and the decoder were in a “siamese” fashion. A siamese network can be seen as two identical subnetworks with shared weights [20]. Each network processes its own input sample and the weights are updated according to a contrastive loss function, so that a model learns to judge whether the inputs belong to the same class or not. For our model the goal is to translate the visual similarity measure between a pair of input images *I*_*1*_, *I*_*2*_ into the mutual distance between a pair of corresponding vector representations *z*_*1*_, *z*_*2*_ in n-dimensional latent space. Moreover, the vector clusters should be compact for each of the image categories and also should be uniformly spread across the latent space to prevent the occurrence of large “blank gaps” which would affect the quality of reconstructed images *I*_*1r*_, *I*_*2r*_. Considering all the specified features, we proposed a loss function as a weighted sum of three components: distance loss, angle loss and pixel loss. In this work after some parameter tuning we set the weights as *w*_*d*_=1, *w*_*a*_=4, *w*_*p*_=2.

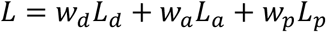

Distance loss function was used to control the mutual distance between latent space representations and was calculated as following:

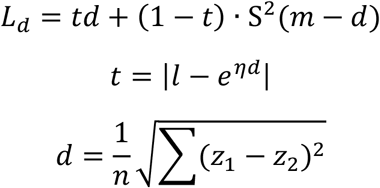

where *l*=1 if the images belong to the same category, otherwise *l*=0. Therefore, target coefficient *t* is close to zero for similar images of the same category;

***S*** is sigmoid function:

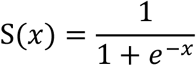

*η* is a distance weighting parameter (in this research we used *η* = 10^−3^);

*m* is margin that separates clusters in latent space. Here we used *m*=1.

In angle loss function cosine similarity metric was used to maintain the uniformness of cluster positions across latent space:

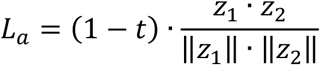

This loss function prevents the category clusters from forming a linear distribution in the latent space, making them form a kind of polygon instead. The mutual distances between the cluster centroids in such distribution is more or less similar, and no a priori preferences are given to any class.

Pixel loss is a common loss function for generative model loss that controls the quality of image reconstruction. For a pair of images, it is a sum of mean square errors for both image reconstructions:

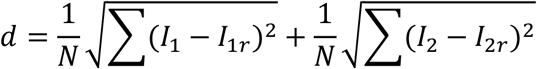

Where N is number of pixels in the images. Another useful feature of a pixel loss is that it makes similar images from the same video clip (e.g. faces of the same person) to have close locations in latent space, therefore the stimuli data tends to form compact subclusters within a common category cluster (see Appendix 1). For our feedback model it means that similar EEG features will be decoded into similar stimuli exemplars (e.g. a particular face) rather than switch chaotically between different exemplars of the category.

The image decoder was trained on a dataset comprised of the image frames taken from the training session video for subject-specific preselected categories. Image pairs were randomly selected to create equal number of same class pairs and different class pairs. Some visualizations for ID model performance can be found in Appendix 1.

### 4.2. EEG feature mapper

The aim of an EEG feature mapping network (FM) is to translate the data from EEG feature domain (*f*) to the image decoder latent space domain (*f*’). Ideally, an image observed by a subject and the EEG recorded at the time of this observation would finally be transformed into the same latent space vector, so that a decoder would produce a proper visualization of what the subject had just seen or imagined:

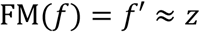

Another problem is to cope with noisy or inconsistent data: while EEG signal properties in real-time recording scenario can significantly vary due to the undetected artifacts or subject getting distracted, the feedback system still should be kept from chaotic image switching as that would put excessive stress on the subject.

The fact that we are working with continuous data gives ground for utilizing the recurrent network models for solving the task. In this research we used long-short term memory (LSTM) cells as recurrent units [21]. We also incorporated the attention mechanism, which makes the model emphasize the key data features and ensures better stability against outliers [22]. A scheme of our EEG feature mapper is shown on Figure 5a and its training routine is presented on Figure 5b.

**Figure 5.**
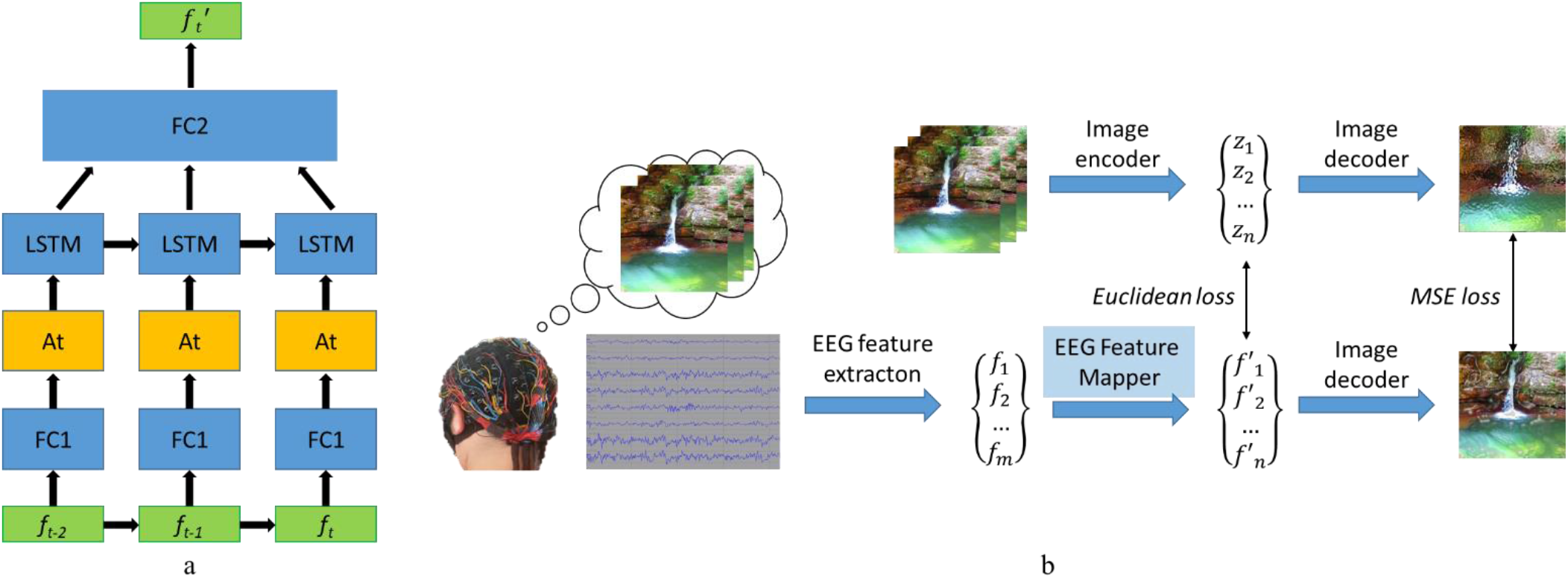
EEG feature mapper. a) Model structure; b) Training routine

A loss function for feature mapper model minimizes the mean square error between EEG and image feature representations both in latent space and in image reconstruction space after decoding:

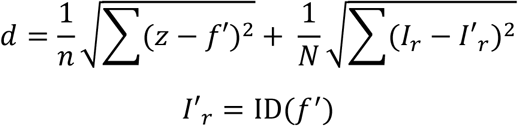

The feature mapper was trained on a dataset comprised of the image frames from the training session video and corresponding 3-second EEG signal windows (centered on the moment of frame onset). EEG feature vectors were extracted using the same method as described in Feature extraction section of this work. Some visualizations for FM network performance can be found in Appendix 1.

### 4.3. Real-time evaluation and results

The proposed neurofeedback model was implemented on Python (with deep learning models implemented using pyTorch library) and run on a machine with an Intel i7 processor, NVIDIA GeForce 1050Ti GPU and 8 Gb RAM. The EEG data was collected in real time via lab streaming layer (LSL) data flow from the amplifier. The processing speed was nearly 3 frames per second, which included incoming EEG data acquisition, filtering, feature extraction and image reconstruction. Before passing to the real-time experiments we extensively tested our neurofeedback model in emulation mode using the data sessions recorded for different subjects. We trained the feedback model using three individual top-recognized categories for each subject. Unfortunately, the objective criteria for feedback visualization quality is yet to be developed, and we had to rely on subjective judgements considering the achieved results. Typically, around 90% the reconstructed imaged were recognizable in terms of category affiliation. The correct correspondence between the classifier predictions and the reconstructed image object category was always established in cases when classifier prediction was more or less stable for a period of 2-3 seconds. This delay time can be regulated by FM network hidden state parameter and can be decreased, which comes at cost of losing the image stability.

The examples of reconstructed images for 4 object categories are shown on Figure 6. More examples can be found in Appendix 1.

**Figure 6.**
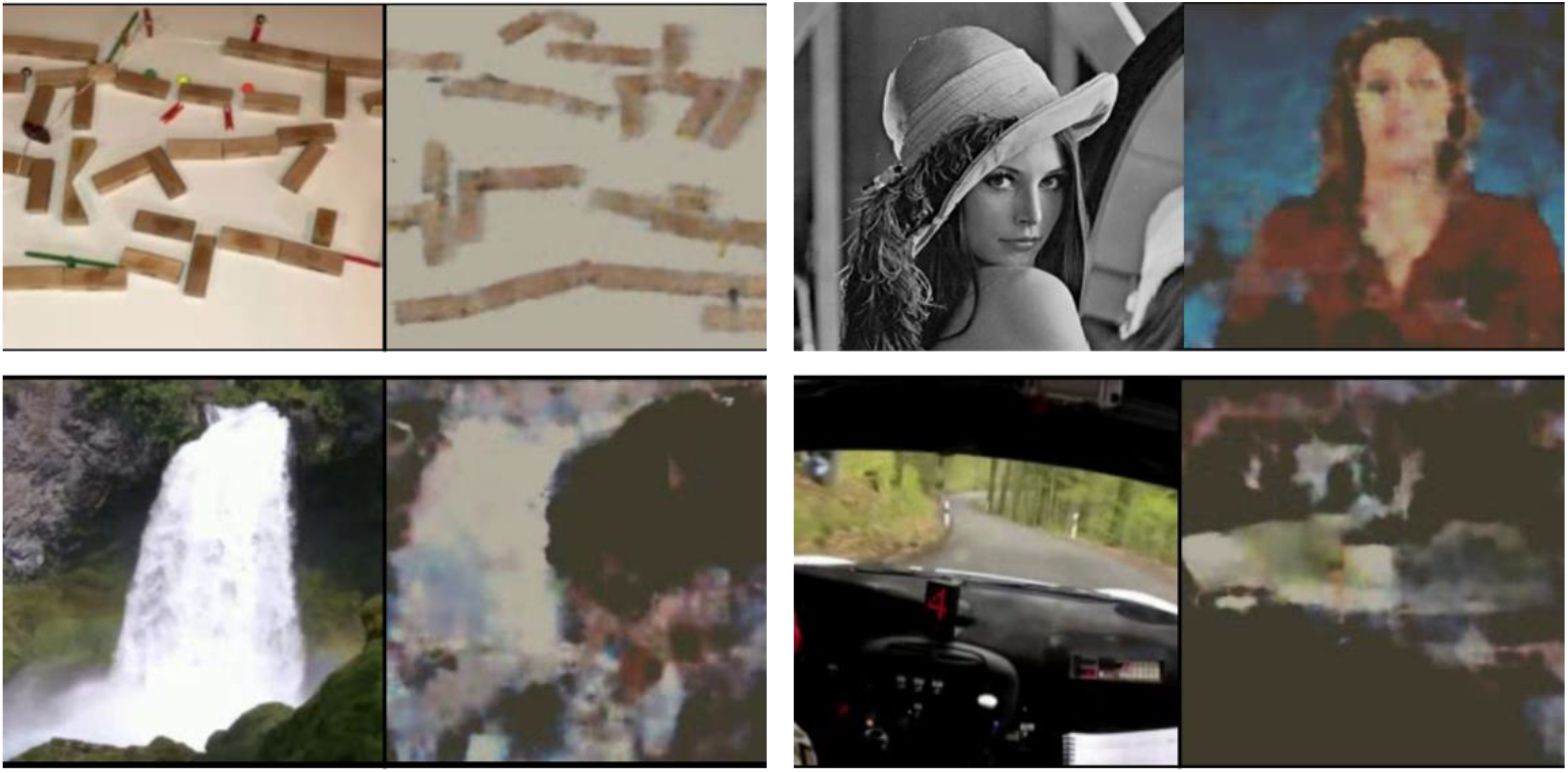
Original images from video stimuli and reconstructed images obtained after processing the co-occurring EEG signal (an original face image replaced by an image sample due to publication policy)

For testing our neurofeedback system in real time we requested one of the subjects to watch the experimental video from the test session once again (Figure 7). The model performance was found to be the same as at the emulation stage. We also requested the subject to watch the presented feedback images instead of the actual video clips and try to switch between the categories at his own will. The subject managed to master imaginary switching between two of three offered categories during 10 minutes of training.

**Figure 7.**
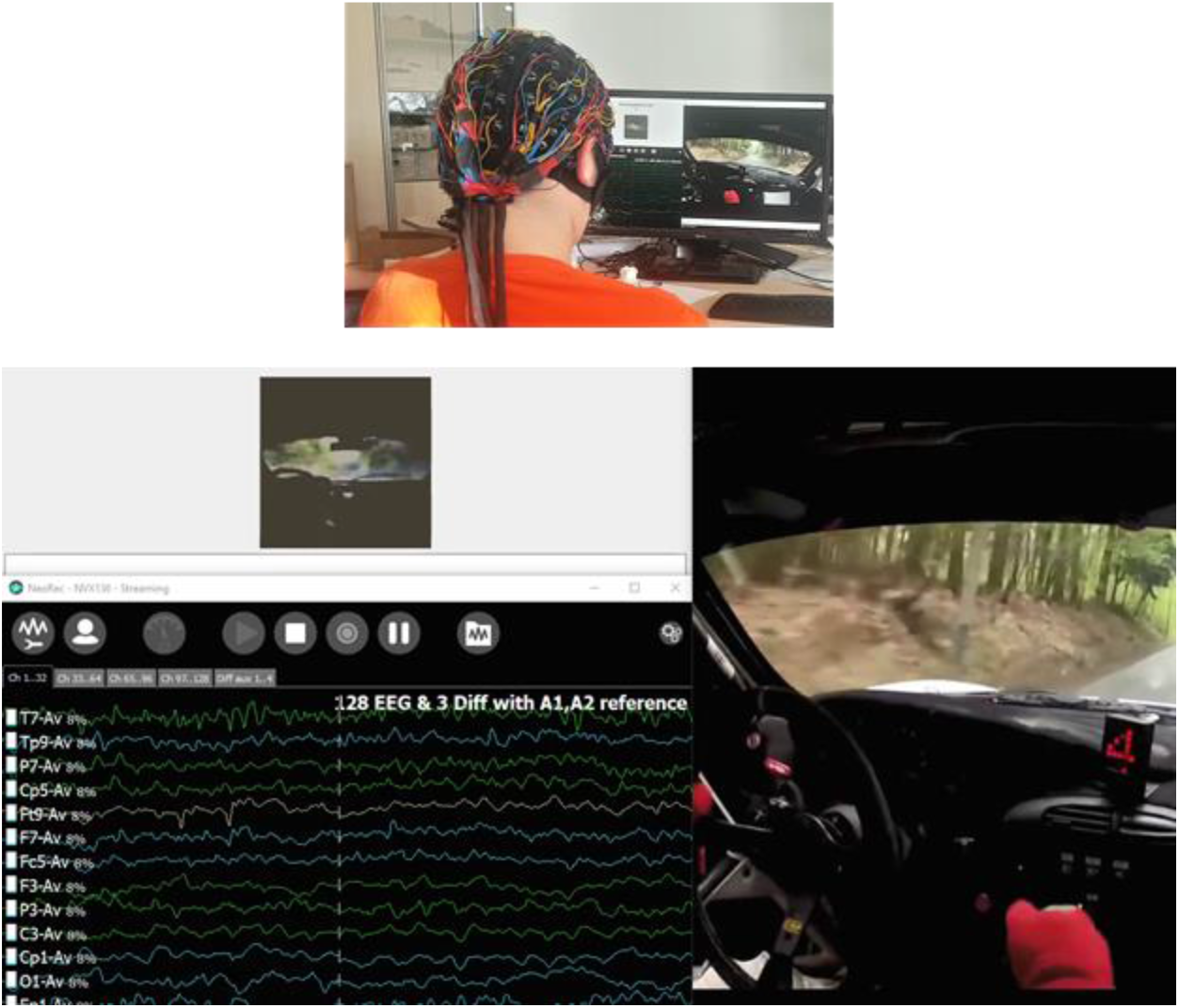
Real-time experimental run. On upper left corner of the screen a reconstructed image of its original (192×192) size can be seen. For this illustration we positioned the EEG recording software window on the same display as the neurofeedback output. During evaluation we hid this window to minimize the distracting factord for the subject.

## 5. Discussion

In this research we explored the effect of continuous visual stimuli presentation on the subject’s brain waves registered in dense noninvasive EEG. The obtained results show that the electrical brain activity can be modulated by presenting subjects with the visual stimuli in the form of video clips. Different classes of objects present in the video clips caused different effect on the brain electric potentials, making it possible to distinguish the discrete scalp EEG patterns which corresponded to each of the object classes and were stable in time. We showed that it was possible to vary the stimuli within each category without affecting the inter-category separability. This makes the proposed method suitable for non-ERP based synchronous BCI implementations.

Our stimulation protocol can be considered as a cognitive test that aims to extract subject-specific stable EEG patterns. Each person has specific reactions to different kinds of visual stimuli, thus, for an effective BCI paradigm a preliminary step of individual stimuli set selection from some excessive basic set could be of a great value. Another benefit of this approach is that no additional cognitive task for attention or memory is required, and the subject can remain completely passive throughout the session. This makes it possible to implement this protocol for the patients with cognitive disorders.

Basing on the results achieved through the developed experimental protocol, we proposed a novel closed-loop BCI system, which is capable of real-time image reconstruction from the subject’s EEG features. We developed a deep learning model which consists of two separately trained deep learning networks, one of which is used for decoding of different categories of images, and the second one transforms the EEG features into the image decoder spatial domain. We demonstrated that the proposed technique can potentially be used for training BCI-naïve subjects by replacing the original stimuli with the subject’s mind-driven image reconstruction model. We suggest that using native feedback could produce a strong self-regulating effect and help a BCI operator to master the imagery commands more effectively.

Further extensive research is required to explore the efficiency of the BCI system in real-world applications. In our future work we will focus on following aspects:

- exploring more visual stimuli categories;
- improvement of reconstructed images quality by incorporating discriminator into image decoder network;
- developing a criteria for assessing the quality of images reconstructed from EEG features
- setting up comparative experiments to evaluate the role of adaptive stimuli selection and visual feedback presentation in BCI operator training speed and quality.

## 7. Acknowledgments

The authors would like to thank Dr. Georgiy Ivanitsky and Prof. Alexei Ossadtchi for the useful discussions related to the details of experimental protocol and EEG data processing.

## 9. Data Availability Statement

The raw data supporting the conclusions of this manuscript will be made freely available by the authors. The video of real-time experiment is available on https://www.youtube.com/watch?v=nf-P3b2AnZw&t=3s

## Appendix 1. Visualization of image reconstruction model performance*

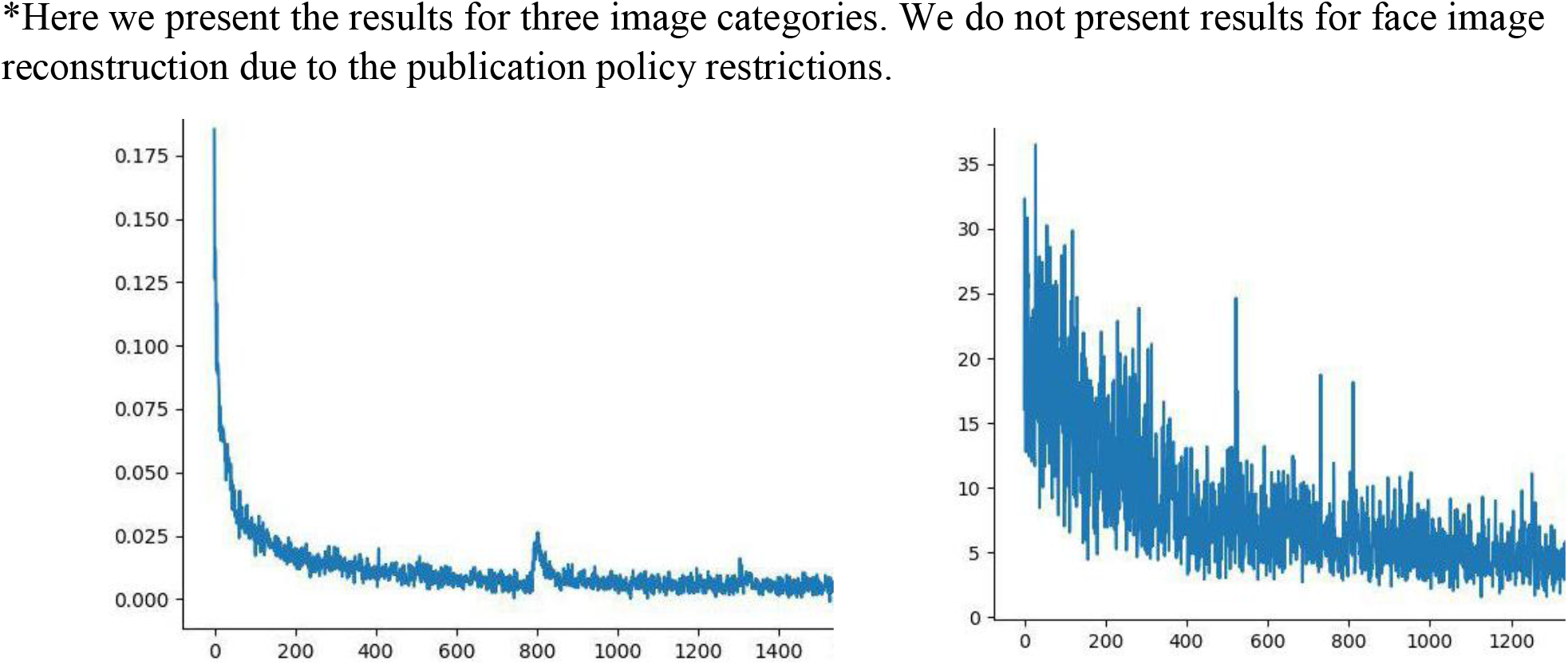
Learning curve of ID (left) and FM (right) models

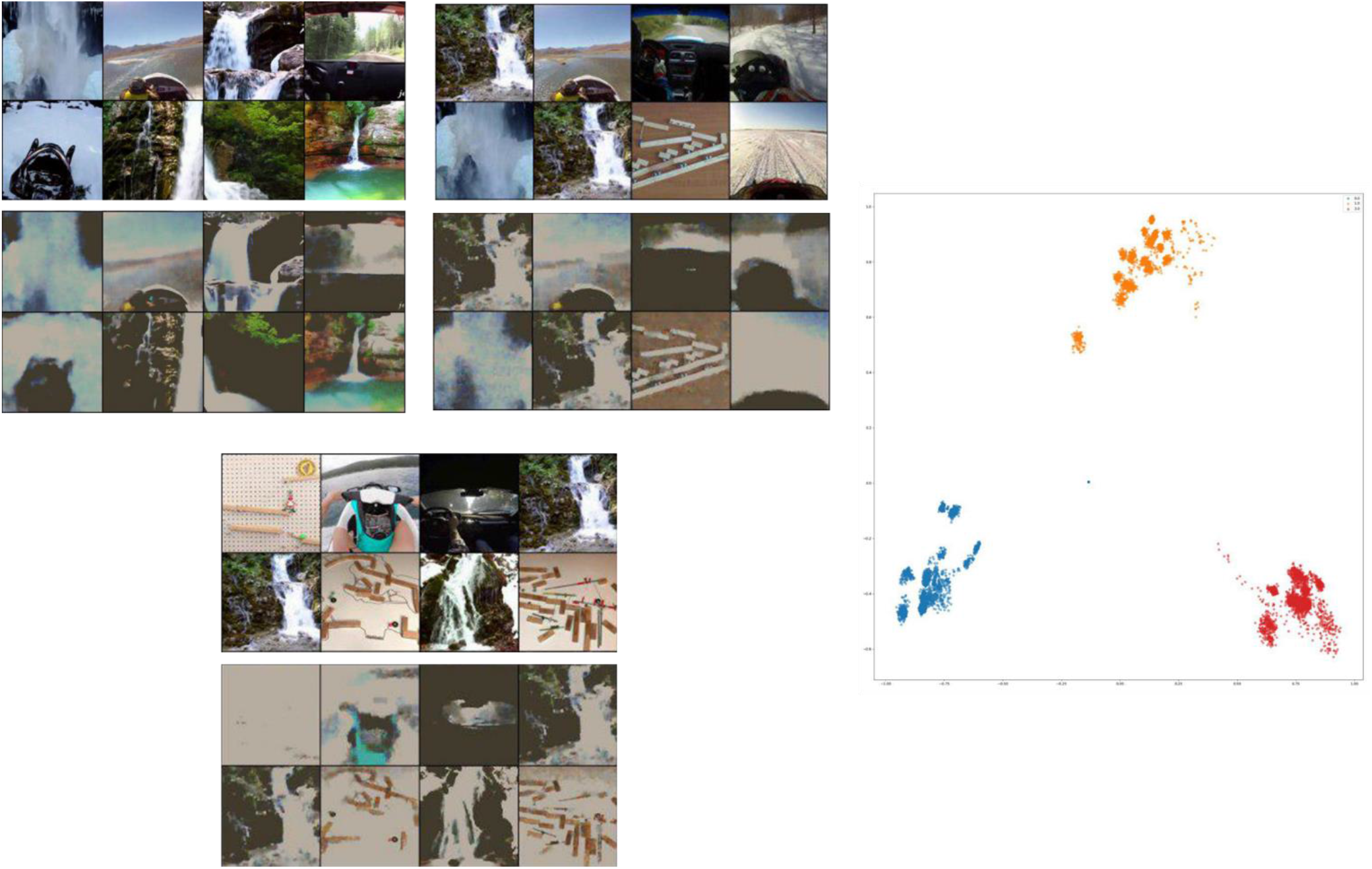
ID performance examples on train data and cluster distributions in the latent space (compared to the corresponding original images from the stimuli video data set).

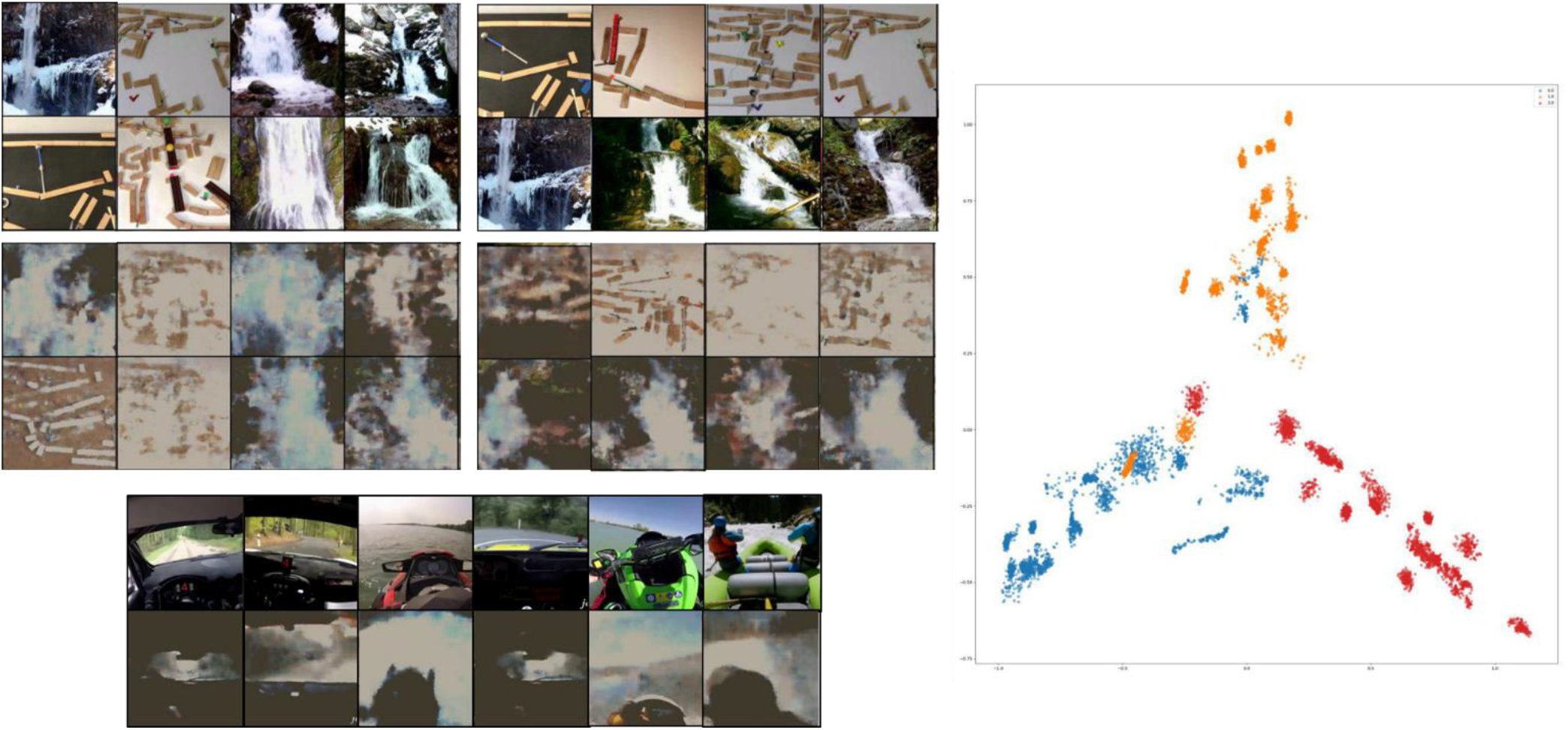
ID performance examples on test data and cluster distributions in the latent space. Note the inner-category subclusters formed by groups of similar images from single clips.

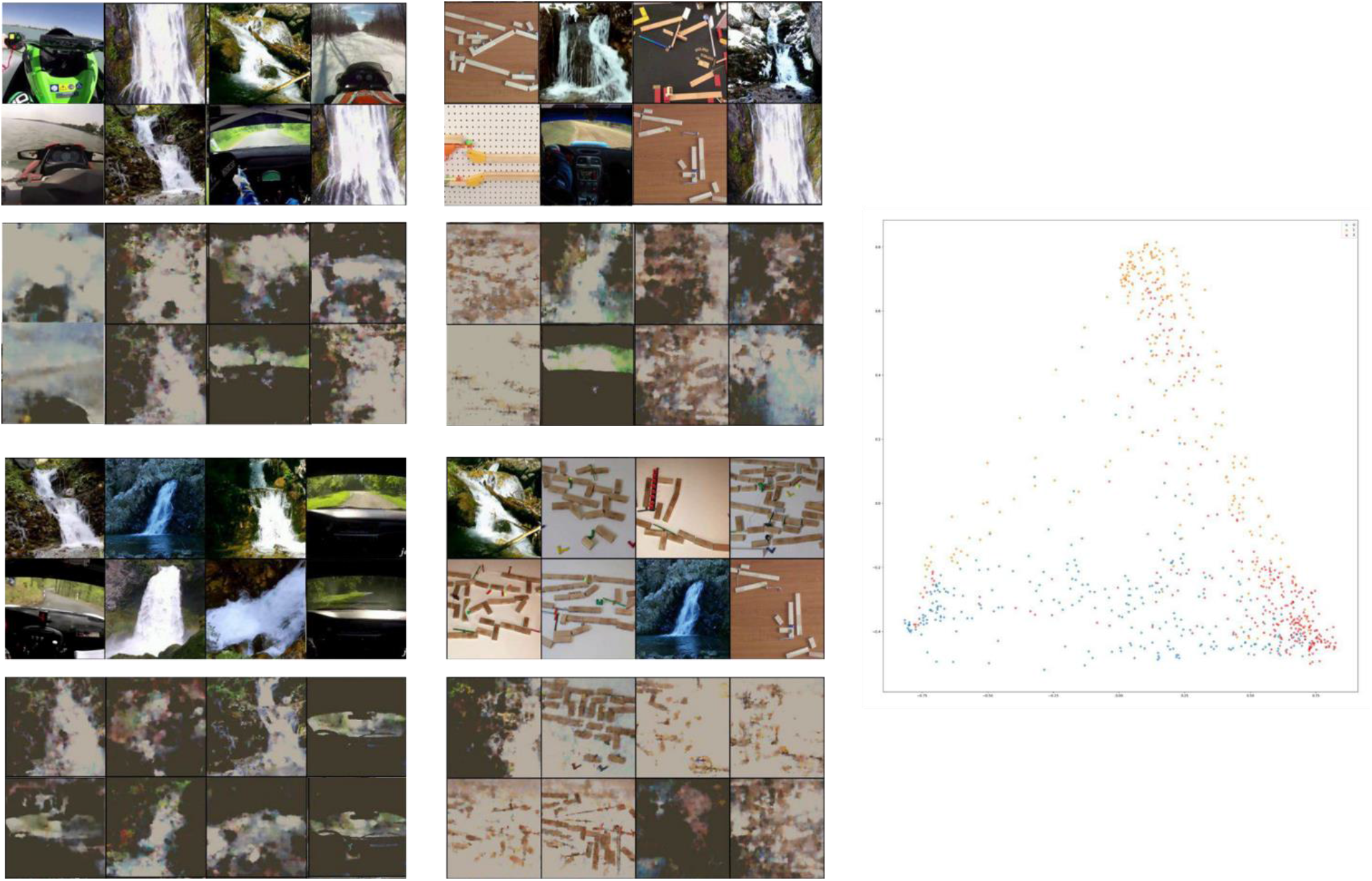
ID performance examples on EEG feature data mapped with FM and the distribution of the mapped data clusters in the latent space.

